# Nutrient Export and Periphyton Biomass in a Stream-Lake Basin from the Patagonian Andean Region

**DOI:** 10.1101/2020.09.07.284893

**Authors:** Alejandro Sosnovsky, Mailén Elizabeth Lallement, Magalí Rechencq, María Valeria Fernández, Eduardo Enrique Zattara, Claudia Silvina Feijoó

**Author notes:** Corresponding author: Alejandro Sosnovsky.

## Abstract

We characterized how land use influenced dissolved nutrients and periphytic algal biomass in an Andean basin from Northwest Patagonia. Nutrient export, especially dissolved inorganic nitrogen increased with human population density. However, no correlation between nutrient concentration and algal biomass was found, which could instead be limited by light availability. Our results suggest that local N-limited ecosystems are liable to eutrophication by increased demographic pressure and that alternative wastewater treatment strategies are necessary for sustainable growth.

---

Physicochemical characteristics of a stream reflect the interactions between water and the basin it flows through. For example, a stream’s N:P ratio depends on water flow paths of each individual watershed (Green *et al.*, 2007). Eutrophication, the enrichment of surface waters with nutrients, is a key human-induced stressor in streams and often a consequence of intensified land use (Dodds & Smith, 2016). Stream eutrophication results in higher export and concentration of phosphorus (P) and nitrogen (N), usually followed by increased algal biomass and Cyanobacteria blooms (Álvarez *et al.*, 2017). Since algal intracellular nutrient concentration and stream nutrient concentration are directly linked, ambient N:P ratio indicate which of these elements could act as a limiting factor to primary productivity of the ecosystem. A review of field studies found that the optimal C:N:P molar ratio of freshwater benthic algae is 158:18:1 (Kahlert, 1998), suggesting that P is limiting when environmental concentration of N:P exceed 18:1, and N is limiting when ratio is < 18:1. Thus, N:P ratio is a simple metric to evaluate the impact of land use changes on nutrient limitation in stream ecosystems.

The ecological characteristics of Northwest Patagonia region are driven by the Andean range. This region has a temperate-cold climate with markedly seasonal precipitation regimes (Paruelo *et al.*, 1998). Its lakes and forests are generally N-limited ecosystems (Díaz *et al.*, 2007; Diehl *et al.*, 2008), while its Andisols-type soils are characterized by high capacity to retain P (Satti *et al.*, 2003). The region harbours a wide network of stream ecosystems subject to a yearly cycle formed by three contrasting hydrological periods: i) stormflow, when chemistry is controlled by lateral flow through the landscape, ii) meltflow, when chemistry is largely influenced by melting snow in the uplands and iii) baseflow, when chemistry is controlled by groundwater inflow (Ahearn *et al.*, 2004; Sosnovsky *et al.*, 2020). Although a considerable fraction of the region has been under an increasing influence from human activities since urban settlement started by the early 1900’s, it is startling that studies linking this influence to eutrophication-driving nutrients like N and P are still scarce (Baffico, 2001; Miserendino *et al.*, 2011; Sosnovsky *et al.*, 2019). We hypothesize that more intensive land use will translate into higher nutrient export and concentration of nutrients in fluvial ecosystems, resulting in increased periphyton biomass on an otherwise nutrient-limited ecosystem. Thus, we aimed to determine during the baseflow period if i) dissolved inorganic nutrient export (i.e., N and P) varied with human population density and ii) changes in nutrient concentration were reflected by changes in periphytic algal biomass.

Research was conducted in a suburban area within the city of San Carlos de Bariloche, Northwest Patagonia, Argentina. This city has approximately 110 000 inhabitants distributed across several drainage basins. One of them, Gutiérrez stream’s drainage basin *(41°09’36.18”S 71°24’37.19”W)*, is large (162 km^2^) and includes a large and deep lake (Gutiérrez Lake, 17 km^2^, 111 m maximum depth) (Fig. 1). Settlements in this basin are not connected to the city’s main sewer system, using instead economic on-site wastewater treatment systems (OWTS) to treat and dispose domestic wastewater. Thus, we can assume that OWTS densities are directly related to local population density within Gutiérrez’s basin. Wastewater undergoes primary treatment in a septic tank via sedimentation and anaerobic digestion in OWTS. Secondary treatment of the septic tank effluents occurs within a rubble bed. This bed has a large aerobic zone, favouring nitrification processes over denitrification ones. The lake of this basin has 7 inflow perennial streams; its seasonal average concentrations of soluble reactive phosphorus (SRP) and dissolved inorganic nitrogen (DIN) are 3.4 mg m^−3^ and 4.6 mg m^−3^, respectively (Díaz *et al.*, 2007). Gutiérrez stream, the outflow of this lake, flows 9 km through a wide gently sloped valley and has a single perennial affluent, the Cascada stream. Gutiérrez’s riparian zone is extensively colonized by the exotic crack willow, *Salix fragilis*, and runs through a series of populated areas. In addition, a trout farm with a total production of ~10 tons year^−1^ pours its sewage into Gutiérrez stream.

**Fig. 1.**
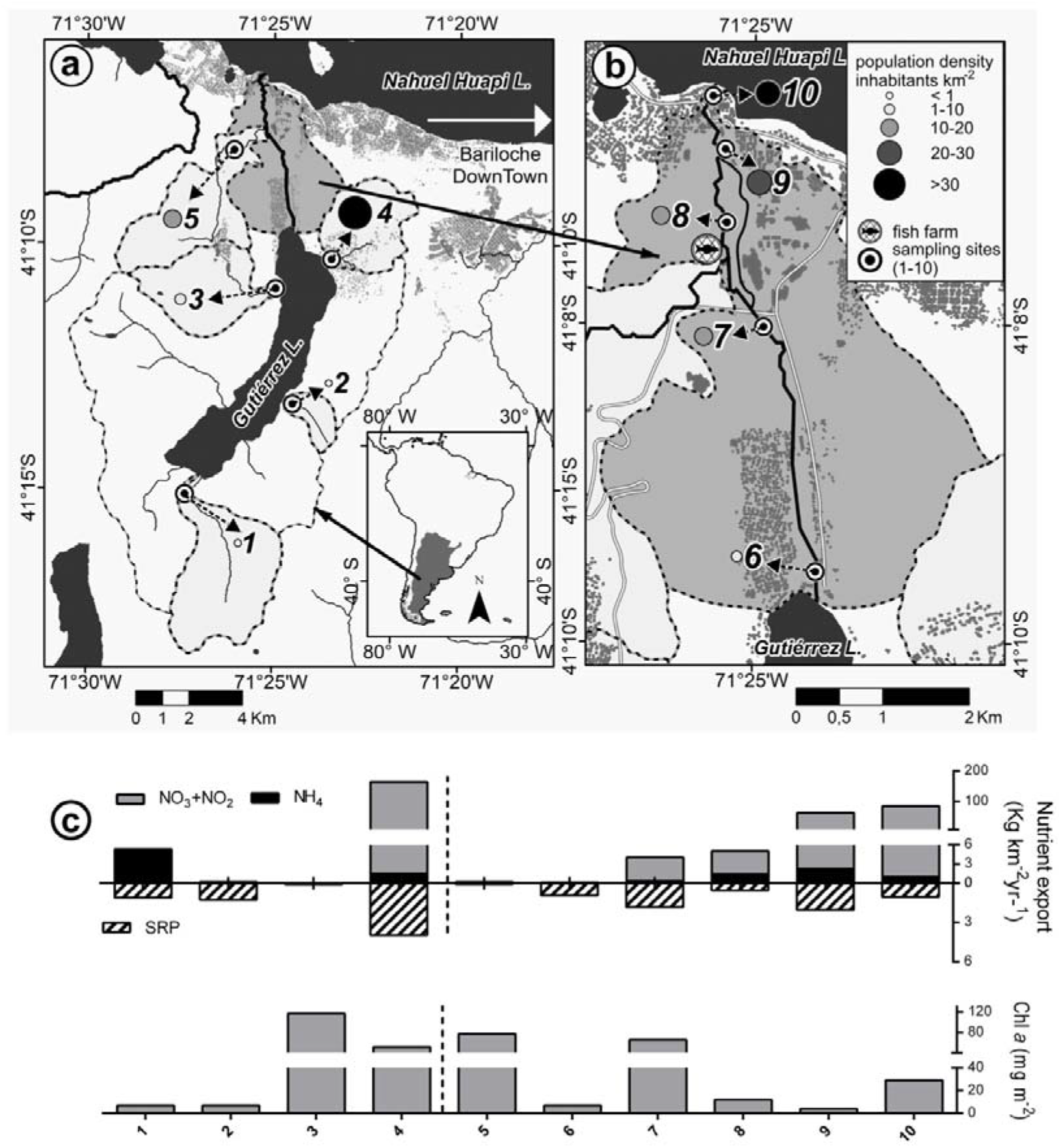
a) Gutiérrez stream basin with the sampled sub-basins highlighted Sampling sites at tributaries and mainstem of Gutiérrez stream are shown: 1) Torrontegui stream, 2) Melgarejo stream. 3) Pescadero stream. 4) Loncochinoco stream. 5) Cascada stream, b) 6-10) Gutiérrez stream. Population zones are shown in grey, c) Export of nutrients and concentration of Chlorophyll *a* (Chi *a*). The vertical dotted line represents the lake, to its left are its tributary streams, to its right are the Gutiérrez stream and its tributary. The Y axes corresponding to dissolved inorganic nitrogen (NO_3_ + NO_2_ + NH_4_) and Chi a have two segments for a better visualization of the data

We surveyed water quality along the Gutiérrez stream basin during year 2016’s baseflow period. Demographic density (inhabitants· km^−2^) was calculated from 2010’s National Census data (https://www.indec.gob.ar). Water samples were taken at four of the headwater lake’s tributary streams (1-4), at the Cascada stream (5) and at five points along the main stream (6-10; Table 1, Fig. 1), measured *in-situ* discharge (Q), electrical conductivity (CE), temperature (T) and turbidity, and estimated canopy cover from riparian vegetation using a concave gridded mirror. We filtered 0.5 liters of water through a MGF Munktell filter to determine the concentration of DIN and SRP. We calculated DIN as the sum of ammonium (NH_4_), nitrates (NO_3_) and nitrites (NO_2_). We analysed NH_4_ by the phenol method, NO_3_ + NO_2_ with a FUTURA Autoanalyzer (Alliance Instruments, Frepillon, France) through a reaction with sulfanilamide with a previous Cu-Cd-reduction for NO_3_ and SRP by the ascorbic acid method (A.P.H.A., 2005). The export nutrients were calculated by the following formula:

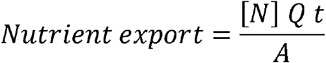

**Table 1.** Morphometric and limnological characteristics of the streams (Str.). Electrical Conductivity (EC); Ammonium (Nh_4_); Nitrates + Nitrites (NO_3_ + NO_2_); Soluble reactive phosphorus (SRP); Dissolved inorganic nitrogen (DIN), Phosphorus:Nitrogen ratio (N:P). Nd: not detectable

| Number/Site | Drainage basin (km<sup>2</sup>) | Discharge (m<sup>3</sup> s<sup>-1</sup>) | EC (μS cm<sup>-1</sup>) | Temperature (°C) | Turbidity (NTU) | Nutrient concentration (mg m<sup>-3</sup>) |  |  |  | N:P (molar) | Canopy (%) |
| --- | --- | --- | --- | --- | --- | --- | --- | --- | --- | --- | --- |
|  |  |  |  |  |  | Nh<sub>4</sub> | NO<sub>2</sub> + NO<sub>3</sub> | DIN | SRP |  |  |
| 1) Str. Torrontegui | 17.0 | 0.158 | 92 | 6.7 | 0.32 | 17.1 | 0.9 | 18.0 | 3.7 | 8.1 | 7 |
| 2) Str. Melgarejo | 2.6 | 0.016 | 198 | 8.1 | 1.47 | nd | 1.1 | 1.1 | 6.5 | 0.2 | 0 |
| 3) Str. Pescadero | 11.3 | 0.006 | 74 | 9.6 | 0.41 | nd | 1.7 | 1.7 | 4.6 | 0.2 | 66 |
| 4) Str. Loncochinoco | 9.4 | 0.099 | 146 | 9.3 | 1.14 | 4.4 | 488.2 | 492.6 | 12.0 | 21.0 | 0 |
| 5) Str. Cascada | 13.0 | 0.003 | 87 | 7.6 | 0.07 | nd | 27.9 | 27.9 | 8.3 | 1.7 | 15 |
| 6) Gutiérrez rise | 133.1 | 1.051 | 70 | 14.8 | 0.17 | nd | 0.5 | 0.5 | 3.7 | 0.1 | 3 |
| 7) Gutiérrez | 144.2 | 1.132 | 76 | 12.6 | 0.24 | 1.0 | 14.8 | 15.8 | 7.4 | 1.3 | 0 |
| 8) Gutiérrez | 158.3 | 0.231 | 88 | 11.5 | 0.83 | 30.2 | 78.3 | 108.5 | 12.0 | 7.6 | 100 |
| 9) Gutiérrez | 160.9 | 0.938 | 80 | 11.7 | 0.32 | 12.0 | 326.0 | 338.1 | 11.0 | 16.6 | 44 |
| 10) Gutiérrez mouth | 161.6 | 0.937 | 80 | 11.7 | 0.35 | 5.3 | 452.3 | 457.6 | 5.5 | 42.4 | 0 |

Where [*N*] is nutrient concentration, Q is discharge, t is time and A is drainage basin area; nutrient export was standardized to kg of nutrient per km^2^ and year (kg km^−2^ yr^−1^).

We also determined chlorophyll *a* (Chl *a*) concentration, corrected for pheopigments (Lorenzen, 1967), as a proxy for periphytic algal biomass. For periphyton sampling, two representative boulders were chosen and scraped, using a piece of cylindrical plastic pipe measuring 16 cm^2^ in cross-section and a brush to standardize the sampled area and remove the algae. Relationships between i) population density and nutrient export, and ii) nutrient concentration and Chl *a* concentration were modelled using standard linear regression, log-transforming variables that failed tests for normal distribution.

Tributary streams to Gutiérrez Lake and stream show a highly heterogeneous distribution of variable values (Fig. 1a, Table 1, supplementary table ST1). These streams had considerably lower temperature and discharge values relative to those found along stream Gutiérrez. In contrast, their nutrient concentrations were higher than those measured at the stream’s headwater (site 6) (Table 1). Torrontegui (site 1) and Melgarejo (site 2) streams, draining sparsely populated basins, showed the lowest concentration of NO_3_ + NO_2_ and Chl *a* (Fig. 1a, Table 1). Despite the low nutrient export at Gutiérrez stream headwater, we detected an increasing trend in DIN export and N:P ratio towards the stream mouth (Fig. 1c, Table 1). This trend was mainly driven by increased NO_3_ + NO_2_ and not NH_4_; ammonium export was highest at the two sites immediately downstream of the trout farm. The increase in DIN in stream G was not accompanied by increases in either SRP exports or Chl *a* concentration (Fig. 1c, Table 1). Instead, Chl *a* concentration was low in sites shaded by a dense riparian cover (sites 8 and 9, Figure 1c, Table 1).

Integrating our results across the whole study basin, we observed that the stream ecosystem responded to an increase in land use intensity, given that DIN and SRP exports were higher at sites draining more densely populated areas (Fig. 2a-b). This relationship was more noticeable for DIN: the linear regression between DIN and population density was significantly higher (*P* = 0.002. compared to *P* = 0.018 for SRP) and explained a larger amount of variance (R^2^ = 0.72, compared to R^2^ = 0.52 for SRP). Similarly, the increase in population density was reflected in an increase in the N:P ratio, which surpasses 18: 1 once population reaches ~ 35 inhabitants per km^2^ (Fig. 2c). This potential effect of land use on nutrient load was not followed by a response from periphytic algal biomass (DIN-Chl *a* R^2^ = 0.02, n = 10, F = 0.17, *P* = 0.69; SRP-Chl *a* R^2^ = 0.01, n = 10, F = 0.05, *P* = 0.82).

**Fig. 2.**
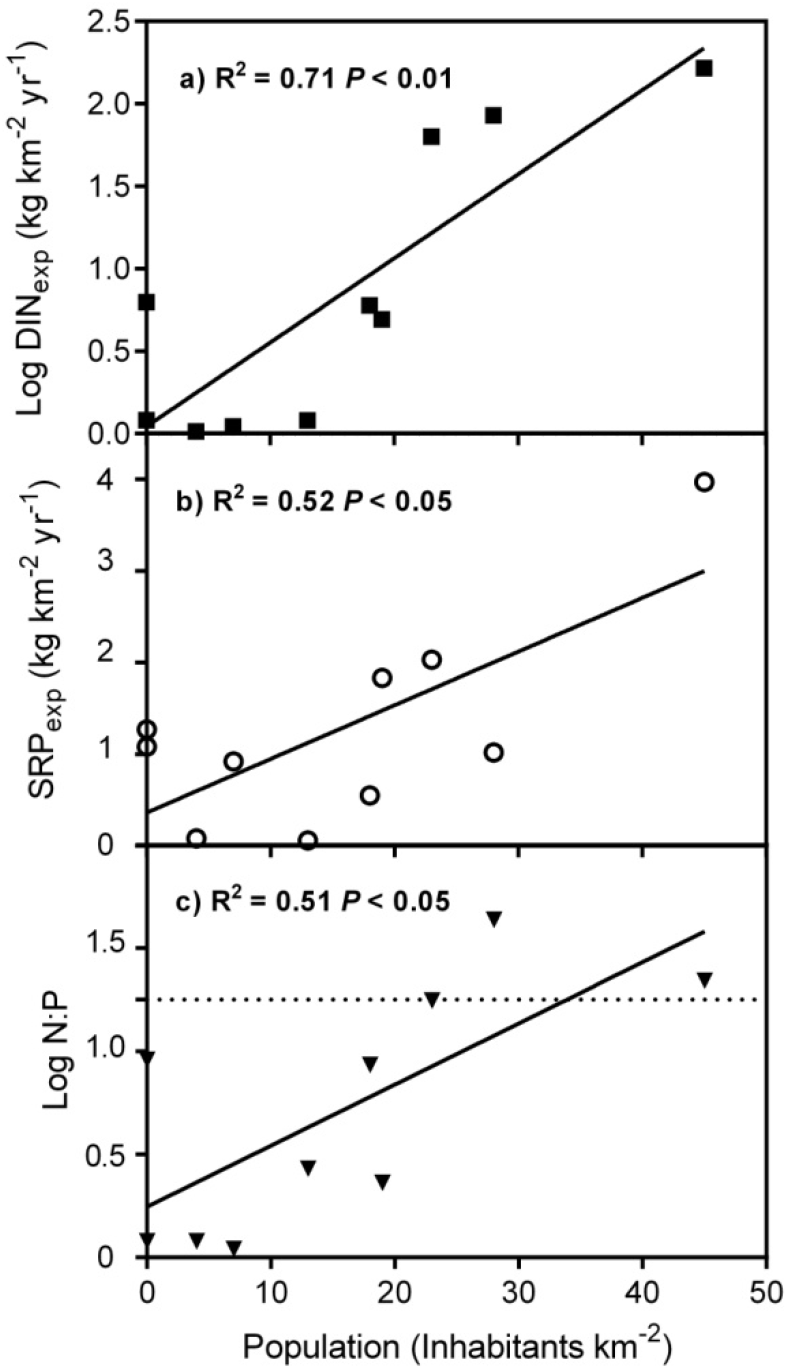
Linear regressions Human density - Nutrients export, a) Dissolved inorganic nitrogen export (DIN_exp_) b) Soluble reactive phosphorus export (SRP_exp_) c) Nitrogen Phosphorus ratio in a moiar basis. Dotted line indicates N:P = 18:1.

Our results reflect a positive relationship between population density and exports of DIN and SRP. On the one hand, the steepest response shown by DIN values indicates that nitrogen was more readily incorporated into the stream ecosystem. This may be partially explained by leeching of nitrates from OWTS towards groundwater (Beal *et al.*, 2005; De & Toor, 2017). In contrast to this pervasive and diffuse source of nitrates to the stream, ammonium increase is better explained by single point sources, likely excretion from fishes at the trout farm. On the other hand, OWTS and Andisol soils seem to be better at retaining phosphorus derived from human activities. Thus, the specific interactions between human settlement, waste treatment strategies and natural environmental characteristics in this area skew nutrient export ratios towards nitrogen enrichment. In mountain areas as the study region, this enrichment becomes most relevant during the baseflow, when groundwater flow path makes the largest contribution to total stream discharge (Ahearn *et al.*, 2004).

The range of ambient N:P ratio has been used to define the transition between N and P limitation in benthic algae (Schanz & Juon, 1983). If the N:P ratio is greater than 20 then P can be assumed to be in limiting supply. If N:P is less than 10 then N can be assumed to be in limiting supply. The distinction of the limiting nutrient when ambient N:P ratios are between 10 and 20 is not precise. The N:P ratio recorded on our study spanned a wide range, from <1 to 42. Thus, the observed lack of response from periphytic algal biomass to the increase in nutrient concentration could be because the limiting nutrient differed between sites, being phosphorous, nitrogen or a colimitation by both them. However, N:P ratio as an indicator of nutrient limitation should be viewed with caution because nutrient requirements for different algal species can vary widely (Allan & Castillo, 2007). In addition, stream ecosystems found within a forest matrix can be limited by light availability (Hill *et al.*, 2001); indeed, we found that sites with low Chl *a* concentration despite high nutrient loads were usually shadier. Thus, we can postulate that in stream ecosystems as the one we studied, the response of primary producers to increased nutrient availability is modulated by the nature and structure of the riparian vegetation. Recently, a significant fraction of the invasive *S. fragilis* population has been removed presenting a great opportunity to test this hypothesis; and more generally, to measure the impact of riparian vegetation on stream productivity.

This work presents some points which deserve further study. NH_4_ concentration in Torrontegui stream was unusually high and merits deeper investigation. Moreover, measurements were limited to the baseflow period, which might have different dynamics relating population density and nutrient exports than the stormflow and meltflow periods of the local hydrologic cycle. Furthermore, our understanding about how periphytic algae respond to nutrient availability would benefit from studying species and chemical composition of these primary producers (Carrillo *et al.*, 2018). Finally, the study of the environmental C:N:P relationships will give us a better comprehension of the auto-heterotrophic dynamics of Andean fluvial ecosystems of Patagonia.

Our research yields insights into stream eutrophication, as it links land uses with physicochemical characteristics of a N-limited drainage basin. The headwater lake acted as an efficient nutrient trap, while residential land use caused increased nutrient load to the basin. Whether this nutrient increase results in algal growth is modulated by the degree of canopy shading. We can conclude that in the Northwest Patagonian region, OWTS as those currently used within Gutiérrez stream basin are likely contributing to eutrophication of streams during the baseflow period. Thus, it would be necessary to consider and evaluate the use of alternative OWTS, with higher denitrification capacity, in order to reduce release of nitrates into groundwater. Furthermore, inhibition of algal response to nitrogen increases by canopy shading could allow build-up of nutrient concentrations to go unnoticed, resulting in potentially harmful blooms if the canopy is removed due to changes in land use. Thus, special care should be taken in preserving the riparian zone of these streams, especially since nitrate concentration of the groundwater might be reduced by denitrification in this zone (Hill, 1996). Given that the Gutiérrez basin is part of a touristic region that can seasonally double its population density, devising mitigation and remediation measurements for human impact is imperative to achieve environmental sustainability.

## Acknowledgements

We acknowledge technician Pablo Alvear for field assistance. Sosnovsky, A., Rechencq, M., Férnández, M.V. and Zattara, E.E. are CONICET researchers. This work was funded by Consejo Nacional de Investigaciones Científicas y Técnicas (CONICET resol. N° 3646/13).

---

**Table 1.** **annexed** Data supporting Figures 1 and 2. Population density and limnological characteristics of the study sites (Str.) Streams. Ammonium (Nh_4_); Nitrates + Nitrites (NO_3_ + NO_2_); Soluble reactive phosphorus (SRP). Nd: not detectable.

| Number/Site | Population<br>(inhabitants km<sup>-2</sup>) | Nutrient export<br>(kg km<sup>-2</sup> y<sup>-1</sup>) |  |  |  | Chlorophyll<br>(mg m<sup>-2</sup>) |
| --- | --- | --- | --- | --- | --- | --- |
|  |  | Nh<sub>4</sub> | NO<sub>2</sub><br>+<br>NO<sub>3</sub> | DIN | SRP |  |
| 1) Str. Torrontegui | 0 | 5.0 | 0.3 | 5.3 | 1.08 | 7 |
| 2) Str. Melgarejo | 0 | nd | 0.2 | 0.2 | 1.27 | 7 |
| 3) Str. Pescadero | 4 | nd | nd | nd | 0.08 | 117 |
| 4) Str. Loncochinoco | 45 | 1.5 | 162.2 | 163.7 | 3.97 | 52 |
| 5) Str. Cascada | 13 | nd | 0.2 | 0.2 | 0.06 | 77 |
| 6) Gutiérrez rise | 7 | nd | 0.1 | 0.1 | 0.92 | 7 |
| 7) Gutiérrez 7 | 19 | 0.3 | 3.7 | 4.0 | 1.83 | 66 |
| 8) Gutiérrez 8 | 18 | 1.4 | 3.6 | 5.0 | 0.55 | 12 |
| 9) Gutiérrez 9 | 23 | 2.2 | 59.9 | 62.1 | 2.03 | 4 |
| 10) Gutiérrez mouth | 28 | 1.0 | 83.1 | 84.1 | 1.02 | 29 |

